# Elevated temperature inhibits SARS-CoV-2 replication in respiratory epithelium independently of the induction of IFN-mediated innate immune defences

**DOI:** 10.1101/2020.12.04.411389

**Authors:** Vanessa Herder, Kieran Dee, Joanna K. Wojtus, Daniel Goldfarb, Christoforos Rozario, Quan Gu, Ruth F. Jarrett, Ilaria Epifano, Andrew Stevenson, Steven McFarlane, Meredith E. Stewart, Agnieszka M. Szemiel, Rute M. Pinto, Andreu Masdefiol Garriga, Sheila V. Graham, Pablo R. Murcia, Chris Boutell

**Affiliations:** MRC-University of Glasgow Centre for Virus Research (CVR), 464 Bearsden Road, Glasgow, G61 1QH, Scotland (UK); University of Glasgow School of Veterinary Medicine, 464 Bearsden Road, Glasgow, G61 1QH, Scotland (UK)

**Keywords:** Respiratory epithelium, SARS-CoV-2, primary human bronchiolar epithelial cells, COVID-19, temperature, immune regulation, RNA-Seq

## Abstract

The pandemic spread of SARS-CoV-2, the etiological agent of COVID-19, represents a significant and ongoing international health crisis. A key symptom of SARS-CoV-2 infection is the onset of fever, with a hyperthermic temperature range of 38 to 41°C. Fever is an evolutionarily conserved host response to microbial infection and inflammation that can influence the outcome of viral pathogenicity and regulation of host innate and adaptive immune responses. However, it remains to be determined what effect elevated temperature has on SARS-CoV-2 tropism and replication. Utilizing a 3D air-liquid interface (ALI) model that closely mimics the natural tissue physiology and cellular tropism of SARS-CoV-2 infection in the respiratory airway, we identify tissue temperature to play an important role in the regulation of SARS-CoV-2 infection. We show that temperature elevation induces wide-spread transcriptome changes that impact upon the regulation of multiple pathways, including epigenetic regulation and lncRNA expression, without disruption of general cellular transcription or the induction of interferon (IFN)-mediated antiviral immune defences. Respiratory tissue incubated at temperatures >37°C remained permissive to SARS-CoV-2 infection but severely restricted the initiation of viral transcription, leading to significantly reduced levels of intraepithelial viral RNA accumulation and apical shedding of infectious virus. To our knowledge, we present the first evidence that febrile temperatures associated with COVID-19 inhibit SARS-CoV-2 replication. Our data identify an important role for temperature elevation in the epithelial restriction of SARS-CoV-2 that occurs independently of the induction of canonical IFN-mediated antiviral immune defences and interferon-stimulated gene (ISG) expression.

## Introduction

The pandemic spread of severe acute respiratory syndrome coronavirus 2 (SARS-CoV-2, SCV2; (1-3)) is an ongoing international health crisis with over 58 million infections and 1.3 million reported deaths worldwide to date (WHO, https://covid19.who.int; November 2020). The spectrum of SCV2 related disease (COVID-19; coronavirus disease 2019) is highly variable, ranging from asymptomatic viral shedding to acute respiratory distress syndrome (ARDS), multi-organ failure, and death. Besides coughing, dyspnoea, and fatigue, fever (also known as pyrexia) is one of the most frequent symptoms of SCV2 infection (4-8). Fever is an evolutionarily conserved host response to microbial infection, which can influence the regulation of host innate and adaptive immune responses (9, 10). Unlike hyperthermia or heat stroke, fever represents a controlled shift in body temperature regulation induced by the expression of exogenous (microbial) and endogenous (host) pyrogenic regulatory factors, including PAMPS (pathogen associated molecular patterns) and pro-inflammatory cytokines (e.g. interleukin 6, IL-6) (9, 10). Body temperature naturally varies throughout the day, with age, sex, and ethnic origin being contributing factors to body temperature regulation (9, 10). In healthy middle-aged adults, fever is defined as a temperature range from 38 to 41°C (ΔT ∼1-4°C above baseline), with low (38 to 39°C), moderate (39.1 to 40°C), high (40.1 to 41.1°C), and hyperpyrexia (>41.1°C) febrile temperature ranges (10). Temperature elevation confers protection against a number of respiratory pathogens (9, 11), with antipyretic drug treatment shown to increase the mortality rate of intensive care unit (ICU) patients infected with influenza A virus (IAV) (12-14).

With respect to COVID-19, up to 90% of hospitalized patients show symptoms of fever (4-8). The majority of patients display low (44%) to moderate (13 to 34%) grade fever (4, 5). COVID-19 ICU patients also show a ∼10% higher prevalence of fever relative to non-ICU patients with milder disease manifestations (6, 15), suggesting fever to play a role during COVID-19 disease progression. *In vitro* studies have shown that SCV2 replicates more efficiently at lower temperatures associated with the upper respiratory airway (33°C), which correlates with an overall weaker interferon (IFN)-mediated antiviral immune response relative to core body temperatures observed in the lower respiratory airway (37°C) (16). These data suggest that tissue temperature could be a significant determinant of SCV2 tropism and immune regulation. However, it remains to be determined what effect temperature elevation above 37°C has on SCV2 infection. We therefore set out to determine the net effect of elevated temperature on SCV2 infection within respiratory epithelial tissue.

Utilizing a three-dimensional (3D) respiratory model that closely mimics the tissue physiology and cellular tropism of SCV2 infection observed in the respiratory airway of COVID-19 patients (16-24), we demonstrate that elevated temperature (≥ 39°C) restricts the replication and propagation of SCV2 in respiratory epithelium independently of the induction of type-I (IFNβ) and type-III (IFNλ) IFN-mediated immune defences. We show that respiratory epithelium remains permissive to SCV2 infection at temperatures up to 40°C, but restricts viral transcription leading to significantly reduced levels of viral RNA (vRNA) accumulation and apical shedding of infectious virus. Importantly, we identify temperature to play an important role in the differential regulation of epithelial host responses to SCV2 infection, including epigenetic, long non-coding RNA (lncRNA), and immunity-related pathways. Collectively, our data identify an important role for tissue temperature in the epithelial restriction of SCV2 replication in respiratory tissue independently of the induction of type-I and type-III IFN-mediated antiviral immune defences.

## Results

### Differentiation of primary human bronchial epithelial cells into ciliated respiratory epithelium supports SARS-CoV-2 replication *in vitro*

In order to establish a respiratory model suitable for studying the effect of temperature on SCV2 replication, we differentiated primary human bronchiolar epithelial (HBE) cells (isolated from a male Caucasian donor aged 63 years) into pseudostratified respiratory epithelium. Haematoxylin and Eosin (H&E) and immunohistochemistry (IHC) staining of respiratory airway cultures demonstrated that these tissues to contain a mixture of epithelial and goblet cells, with significant levels of apical ciliation and expression of ACE2 (Figure 1A), the principal surface receptor for SCV2 and major determinant of tissue tropism (25-28). Infection of respiratory airway cultures with SCV2 (strain England 2; 10^4^ PFU/tissue) at 37°C demonstrated that these tissues support SCV2 infection and replication, with intraepithelial and apical vRNA accumulation detectable by *in situ* hybridization by 96 h post-infection (Figure 1A). Notably, we observed discrete clusters of SCV2 RNA accumulation within the respiratory epithelium, indicative of SCV2 cell-type specific tropism and/or localized patterns of immune restriction (see below; (29, 30)). The overall morphology of the respiratory epithelium remained largely intact, with little to no discernible shedding of the ciliated surface epithelium over the time course of infection (Figure 1A). Measurement of genome copies (RT-qPCR) and infectious virus (TCID_50_) within apical washes collected over time (24 to 144 h) demonstrated the linear phase of virus shedding to occur between 48 and 96 h, with peak titres at 120 h post-infection (Figure 1B to E).

**Figure 1.**
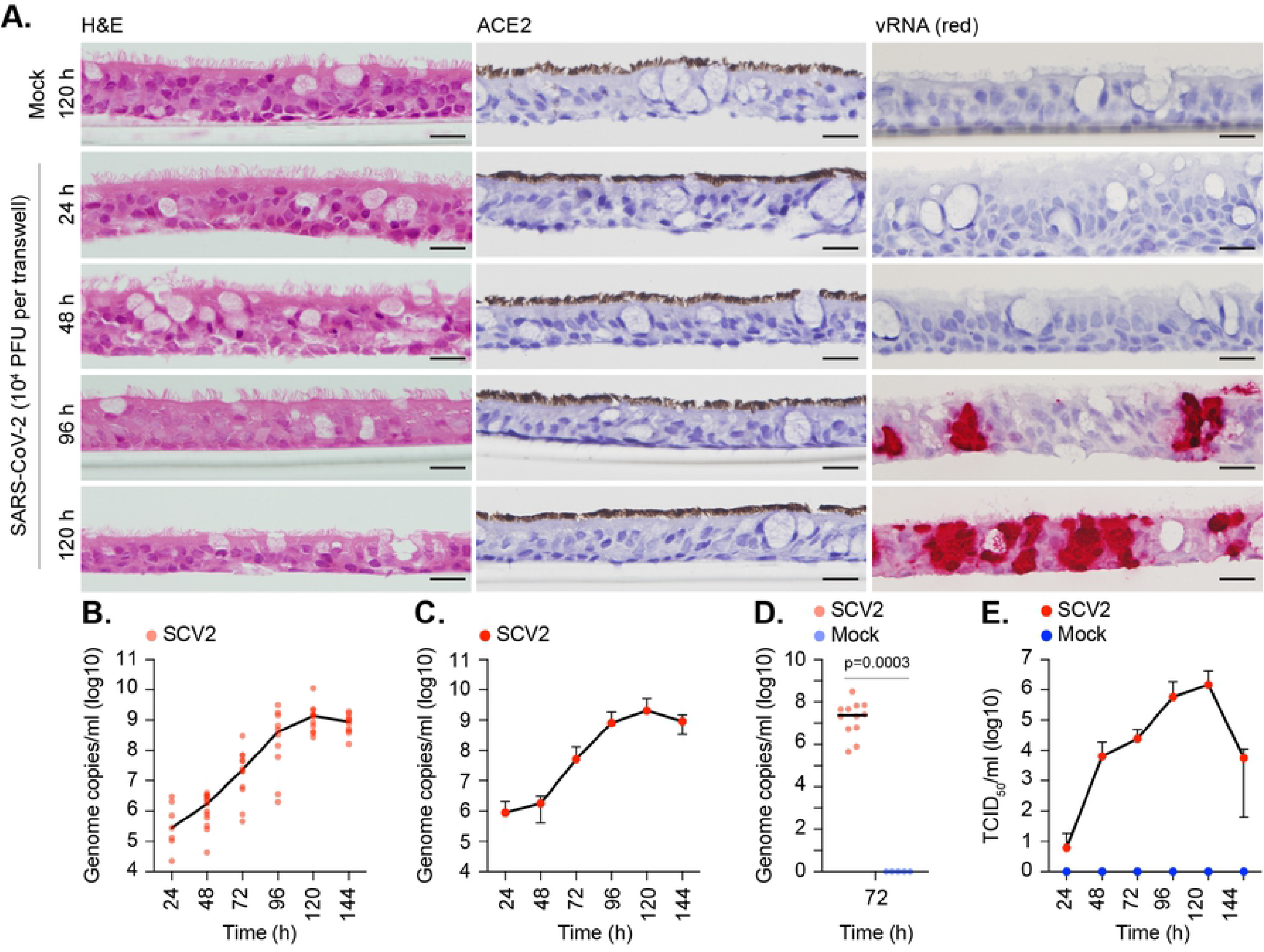
Differentiation of primary bronchial epithelial airway cultures supports SARS-CoV-2 replication. Primary bronchiole epithelial cells were seeded onto 6.5mm transwells and grown to confluency prior to differentiation under air liquid interface (ALI) conditions for ≥35 days. Ciliated respiratory cultures were mock (media only) or SARS-CoV-2 (SCV2; 10^4^ PFU/well) infected at 37°C for the indicated times (hours, h). (A) Representative images of H&E (left-hand panels), ACE2 (brown, middle panels), or detection of SCV2 RNA by *in situ* hybridization (red, right-hand panels) stained sections. Haematoxylin was used as a counter stain. Scale bars = 20 µm. (B/C) Genome copies per ml of SCV2 in apical washes harvested over time as determined RT-qPCR. N≥7 derived from a minimum of three independent biological experiments per sample condition. (B) Black line, median; all data points shown. (C) Means and SD shown. (D) Specificity of RT-qPCR to detect vRNA in apical washes taken from mock (blue circles) or SCV2 infected (red circles) tissues at 72 h (as in B). N≥5 derived from a minimum of three independent biological experiments per sample condition. Black line, median; all data points shown; p=0.0003, Mann-Whitney *U*-test. (E) TCID_50_ assay measuring infectious viral load in apical washes harvested from mock (blue circles) or SCV2 infected (red circles) tissues over time. Means and SD shown. N≥5 derived from a minimum of three independent biological experiments per sample condition.

In order to determine which cellular pathways were modulated in response to SCV2 infection, we performed RNA-Seq analysis on RNA extracted from mock-treated or SCV2-infected respiratory cultures incubated at 37°C for 72 h, a time point in the linear phase of viral shedding (Figure 1C to E). Out of the 787 differentially expressed genes (DEGs) (Figure 2A, Supplemental File S1; p<0.05, ≥ 1.5 or ≤ -1.5 log2 Fold Change [log2 FC]), pathway analysis identified DEG enrichment in immune system and cytokine related pathways to be significantly upregulated in SCV2-infected relative to mocked-treated tissues (Figure 2B, C p<0.0001). Type-I (IFNβ, *IFNB1*) and type-III (IFNλ, *IFNL1-3*) IFNs, along with a subset of interferon stimulated genes (ISGs; e.g. *BST2, ISG15, Mx1*, and *ZBP1* amongst others), and other pro-inflammatory cytokines (*IL6 and IL32*) were upregulated (Figure 2D). Indirect immunofluorescence staining of tissue sections demonstrated that induced ISG expression coincided with productively infected areas of the respiratory epithelium (Figure 2E, Mx1), identifying localized areas of immune regulation within infected tissue. Our findings are consistent with previous reports showing that SCV2 infection induces an IFN-mediated antiviral immune response upon infection of epithelial tissue (16, 18-21, 30-32).

**Figure 2.**
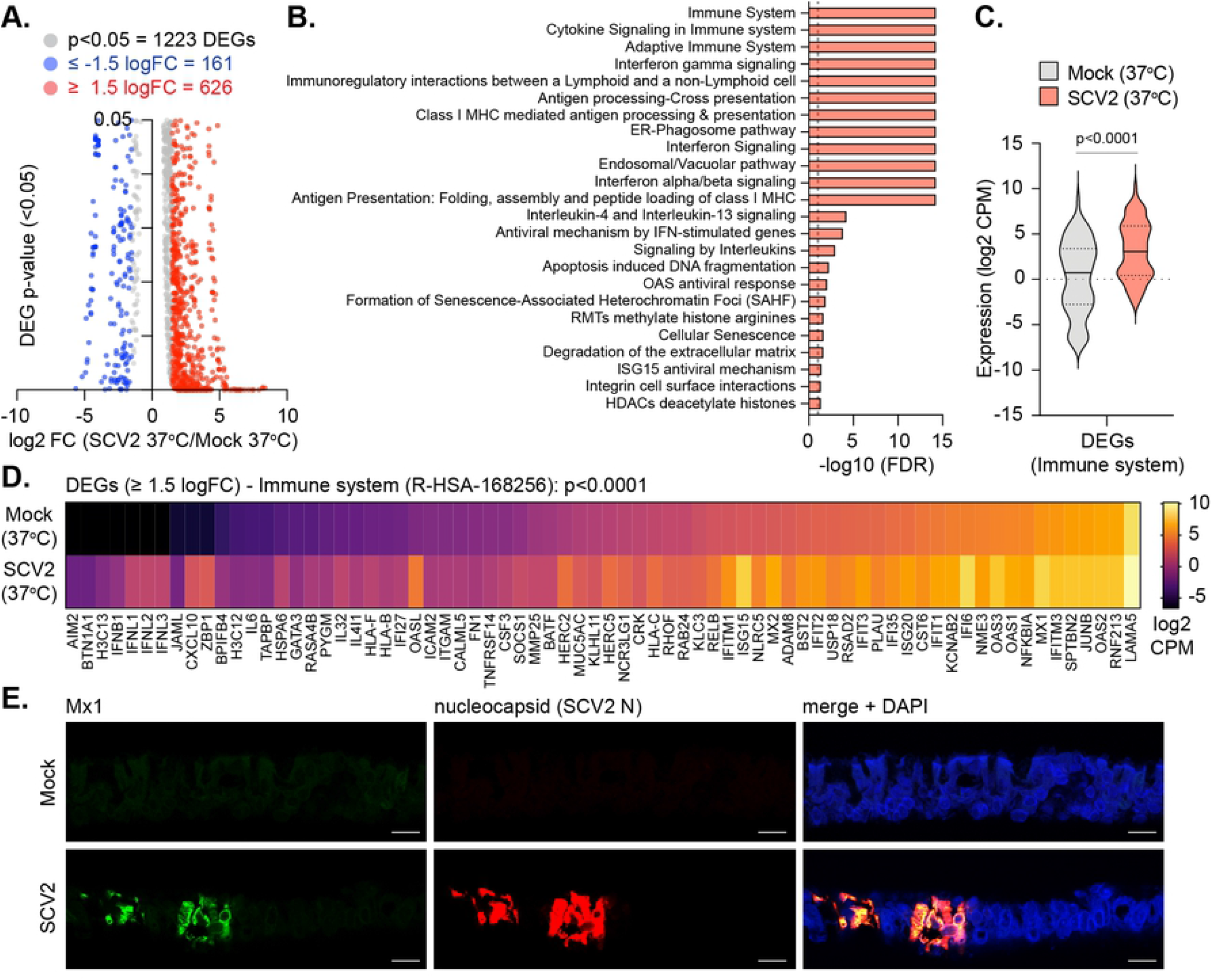
SARS-CoV-2 infection of respiratory airway cultures induces a type-I and type-III IFN response. Ciliated respiratory cultures were mock (media only) or SARS-CoV-2 (SCV2; 10^4^ PFU/well) infected at 37°C for 72 h prior to RNA extraction and RNA-Seq. (A) Scatter plots showing high confidence (p<0.05) differentially expressed gene (DEG) transcripts identified between mock and SCV2 infected cultures (grey circles); DEGs ≥ 1.5 log2 FC, red circles; DEGs ≤ -1.5 log2 FC, blue circles. (B) Reactome pathway analysis of mapped DEGs (p<0.05, ≥ 1.5 log2 FC). Pathways enriched for DEGs with an FDR (corrected over-representation *P* value) < 0.05 (plotted as -log10 FDR) shown (red bars). Dotted line, threshold of significance (-log10 FDR of 0.05). (C) Expression profile (log2 CPM) of Immune system (R-HSA168256) related DEGs (identified in B). Black line, median; dotted lines; 5^th^ and 95^th^ percentile range; p<0.0001, Mann-Whitney *U*-test. (D) Expression levels (log2 CPM) of individual Immune system DEGs (identified in B); p<0.0001, Wilcoxon matched-pairs sign rank test. (A to D) RNA-Seq data derived from RNA isolated from three technical replicates per sample condition. Data analysis is presented in Supplemental File S1. (E) Indirect immunofluorescence staining of tissue sections showing Mx1 (green) and SCV2 nucleocapsid (N, red) epithelial localization. Nuclei were stained with DAPI. Scale bars = 20 µm.

### Respiratory airway cultures induce a heat stress response at elevated temperature

We next investigated the influence of temperature on our respiratory epithelium model. Mock treated respiratory cultures were incubated at 37, 39, or 40°C (representative of core body temperature and low to moderate grade febrile temperatures, respectively) for 72 h prior to tissue fixation or RNA extraction and RNA-Seq. H&E staining demonstrated no obvious morphological changes to the respiratory epithelium upon incubation at elevated temperature (Figure 3A), although an increase in epithelium thickness was detected relative to tissues incubated at 37°C (Figure 3B; 39°C p<0.0001, 40°C p<0.0001). RNA-Seq analysis demonstrated no difference in the expression level of ACE2 (Figure 3C; p=0.4001) or a reference set of genes known to be constitutively expressed across a wide range of cell types and tissues (Figure 3D; 24 genes, p=0.9106; (33-35)). These data indicate that cellular transcription remains largely unperturbed at elevated temperatures up to 40°C. Out of the 650 DEGs identified (Figure 3E, Supplemental File S2; p<0.05, ≥ 1.5 or ≤ -1.5 log2 FC), pathway analysis identified DEG enrichment in multiple pathways in tissues incubated at 40°C relative to 37°C, including meiotic recombination, DNA methylation, rRNA expression, and collagen metabolism (Figure 3F). DEG enrichment was also observed in cellular pathways relating to heat stress (Figure 3F black arrow, 3G; R-HSA-3371556) and cellular response to external stimuli (Figure 3G; R-HSA-8953897). Analysis of complete gene sets associated with the cellular response to heat stress (Figure 3H, p=0.0013) or cellular response to heat (Figure S1, p=0.0018; GO:0034605) identified that these pathways were upregulated in respiratory tissues incubated at 40°C relative to 37°C (Supplemental File S2). We conclude that our respiratory tissue model induces a heat stress response at elevated temperature without visible damage to the epithelium or induction of IFN-mediated antiviral immune defences.

**Figure 3.**
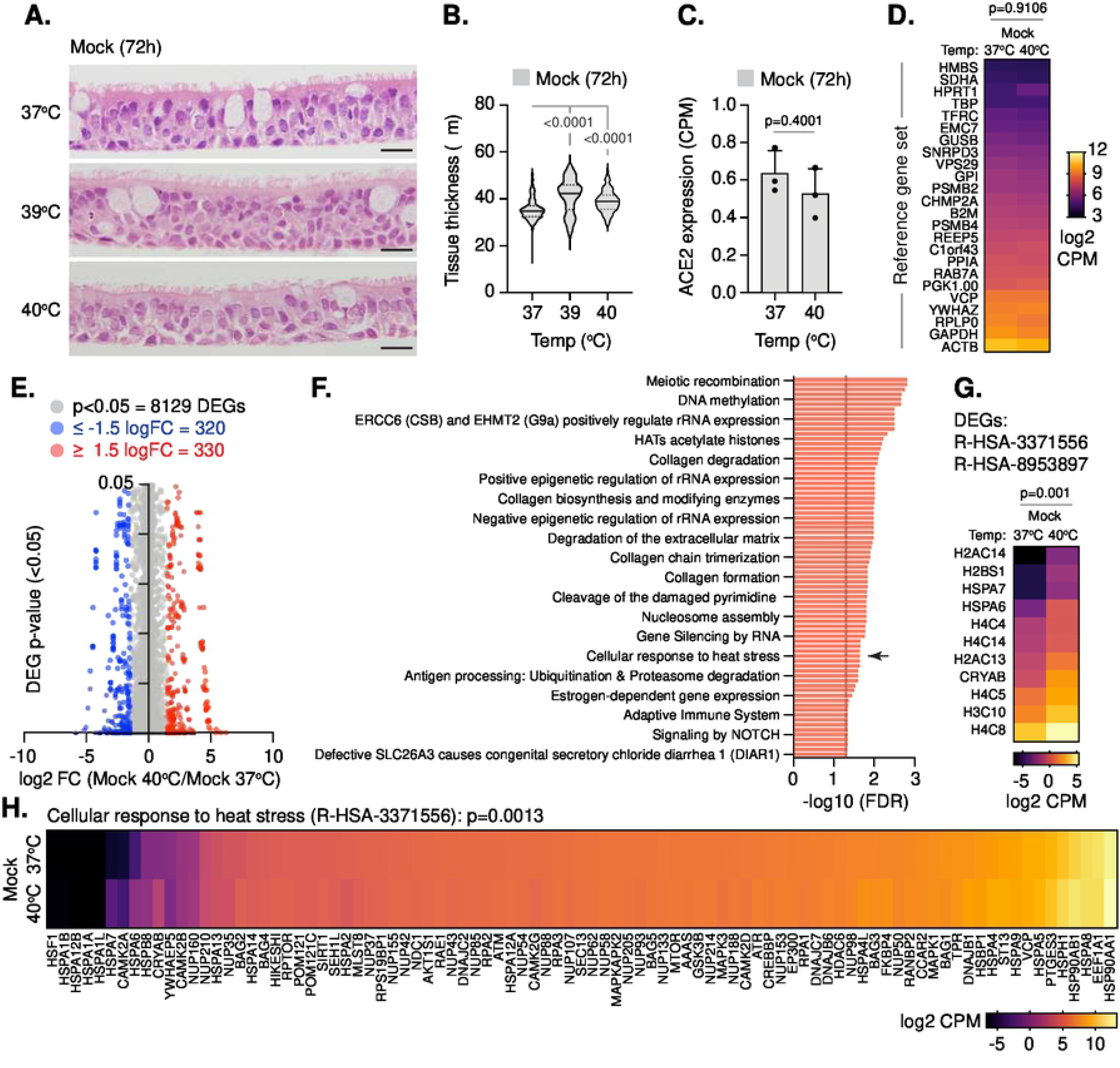
Respiratory airway cultures induce a heat stress response upon incubation at elevated temperature. Ciliated respiratory cultures were incubated at 37, 39, or 40°C for 72 h prior to fixation or RNA extraction and RNA-Seq. (A) Representative images of H&E stained sections. Scale bars = 20 µm. (B) Quantitation of respiratory tissue thickness (μm) at 72 h. N=300 measurements derived from three technical replicates per sample condition. Black line, median; dotted lines; 5^th^ and 95^th^ percentile range; p-values shown, one-way ANOVA Kruskal-Wallis test. (C) ACE2 expression levels (CPM) in respiratory cultures incubated at 37 or 40°C; means and SD shown; p=0.4001, Mann-Whitney *U*-test. (D) Expression values (log2 CPM) of a reference gene set (24 genes; (33-35)) in respiratory cultures incubated at 37 or 40°C; p=0.9106, Mann-Whitney *U*-test. (E) Scatter plots showing high confidence (p<0.05) differentially expressed gene (DEG) transcripts identified between respiratory cultures incubated at 37 or 40°C (grey circles); DEGs ≥ 1.5 log2 FC, red circles; DEGs ≤ -1.5 log2 FC, blue circles. (F) Reactome pathway analysis of mapped DEGs (p<0.05, ≥ 1.5 log2 FC). Pathways enriched for DEGs with an FDR < 0.05 (plotted as -log10 FDR) shown (red bars; every 4^th^ bar labelled). Dotted line, threshold of significance (-log10 FDR of 0.05). (G) Expression values (log2 CPM) of DEGs identified with cellular response to heat stress (R-HSA-3371556; arrow in B) and cellular response to external stimuli (R-HSA-8953897) pathways; p=0.001, Wilcoxon matched-pairs sign ranked test. (H) Expression levels (log2 CPM) for all genes associated with cellular response to heat stress pathway (R-HSA-3371556); p=0.0013, Wilcoxon matched-pairs sign ranked test. (C to H) RNA-Seq data derived from RNA isolated from three technical replicates per sample condition. Data analysis is presented in Supplemental File S2.

### Elevated temperature restricts the replication of SARS-CoV-2 in respiratory airway cultures

We next examined the effect of temperature on SCV2 replication in our 3D tissue model. Respiratory cultures were incubated at 37, 39, or 40°C for 24 h prior to mock treatment or SCV2 infection and continued incubation at their respective temperatures. Measurement of genome copies and infectious virus within apical washes collected over time (24 to 72 h) demonstrated that extracellular SCV2 titres were significantly decreased at both 39 and 40°C relative to 37°C (Figure 4A to D). RT-qPCR analysis of RNA isolated from infected tissues at 72 h post-infection demonstrated significantly lower levels of intracellular vRNA in tissues incubated at 39 or 40°C (Figure 4E; 39°C p=0.0278, 40°C p=0.0033). These data indicate that respiratory tissue remains permissive to SCV2 infection (tissue entry) but refractory to SCV2 replication at elevated temperatures ≥39°C. To our knowledge, these data identify for the first-time a temperature-dependent growth defect in SCV2 replication at elevated temperature (>37°C).

**Figure 4.**
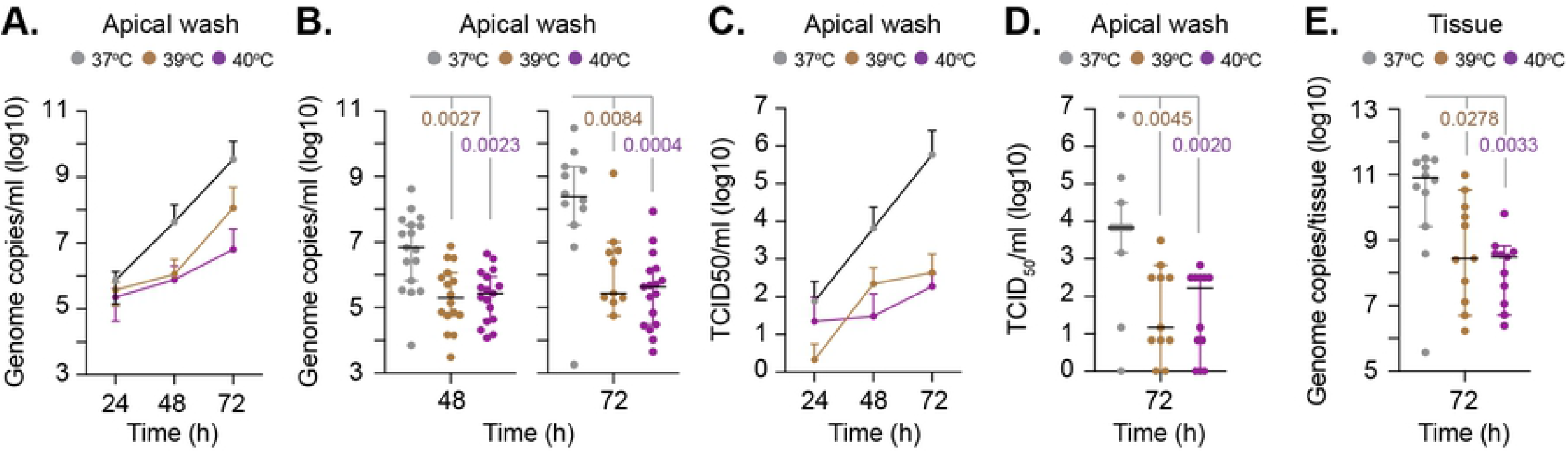
Elevated temperature restricts SARS-CoV-2 replication in respiratory airway cultures. Ciliated respiratory cultures were incubated at 37, 39, or 40°C for 24 h prior to mock (media only) or SARS-CoV-2 (SCV2; 10^4^ PFU/well) infection and incubation at the indicated temperatures. Apical washes were collected over time (as indicated) and tissues harvested at 72 h for RNA extraction and RNA-Seq. (A/B) Genome copies per ml of SCV2 in apical washes harvested over time as determined RT-qPCR. N≥11 derived from a minimum of three independent biological experiments per sample condition. (A) Means and SD shown. (B) Black line, median; whisker, 95% confidence interval; all data points shown; p-values shown, one-way ANOVA Kruskal-Wallis test. (C/D) TCID_50_ assay measuring infectious viral load in apical washes harvested from SCV2 infected respiratory cultures over time. N≥11 derived from a minimum of three independent biological experiments. (C) Means and SD shown. (D) Black line, median; whisker, 95% confidence interval; all data points shown; p-values shown, one-way ANOVA Kruskal-Wallis test. (E) Genome copies per tissue of SCV2 as determined RT-qPCR. N≥11 derived from a minimum of three independent biological experiments per sample condition. Black line, median; whisker, 95% confidence interval; all data points shown; p-values shown, one-way ANOVA Kruskal-Wallis test.

In order examine the underlying mechanism(s) of this intracellular restriction, we compared the relative DEG profiles (p<0.05, ≥ 1.5 or ≤ -1.5 log2 FC) from mock-treated or SCV2-infected tissues at 37 or 40°C (Figure 5, S2, S3, Supplemental Files S1 to S4). Distinct patterns of DEG expression were detected between each paired condition analyzed (Figure 5A, B; Supplemental File S5), with clusters of gene commonality (Figure 5B, purple lines) or shared gene pathway ontology (Figure 5B, blue lines). We also identified lncRNAs and micro-RNAs (miRNAs) to be differentially expressed between temperature and SCV2 infection conditions (Figure S4; Supplemental File S6). Pathway analysis identified DEG enrichment to be highly specific for each paired condition analyzed, with a clear enrichment in immunity related pathways at 37°C relative to all other conditions (Figure 5C). These data indicate that tissue temperature plays an important role in the differential regulation of transcriptional host responses to SCV2 infection of epithelial tissue. To examine the influence of temperature on immune regulation further, we directly compared the transcriptome profiles from SCV2 infected tissues at 37 and 40°C. Out of the 859 DEGs identified (Figure 6A; p<0.05, ≥ 1.5 or ≤ -1.5 log2 FC), pathway analysis identified immune system and cytokine related pathways to be suppressed during SCV2 infection at 40°C relative to 37°C (Figure 6B, C; Supplemental File S7), findings consistent with a failure of SCV2 to induce the expression of type-I (INFβ, *IFNB1*) or type-III (IFNλ, *IFNL1-3*) IFNs at 40°C (Figure 6D). Analysis of this subset of differentially expressed immune genes identified a decrease in gene expression during SCV2 infection at 40°C (mean log2 counts per million [CPM] 1.147, SD 3.656) relative to mock treatment at 37°C (mean log2 CPM 1.578, SD 3.716; Figure 6C). These data suggest that SCV2 infection of respiratory tissue at 40°C is not sufficient to induce a robust innate immune response, indicating that the observed temperature restriction of SCV2 at elevated temperature occurs independently of IFN-mediated antiviral immune defences.

**Figure 5.**
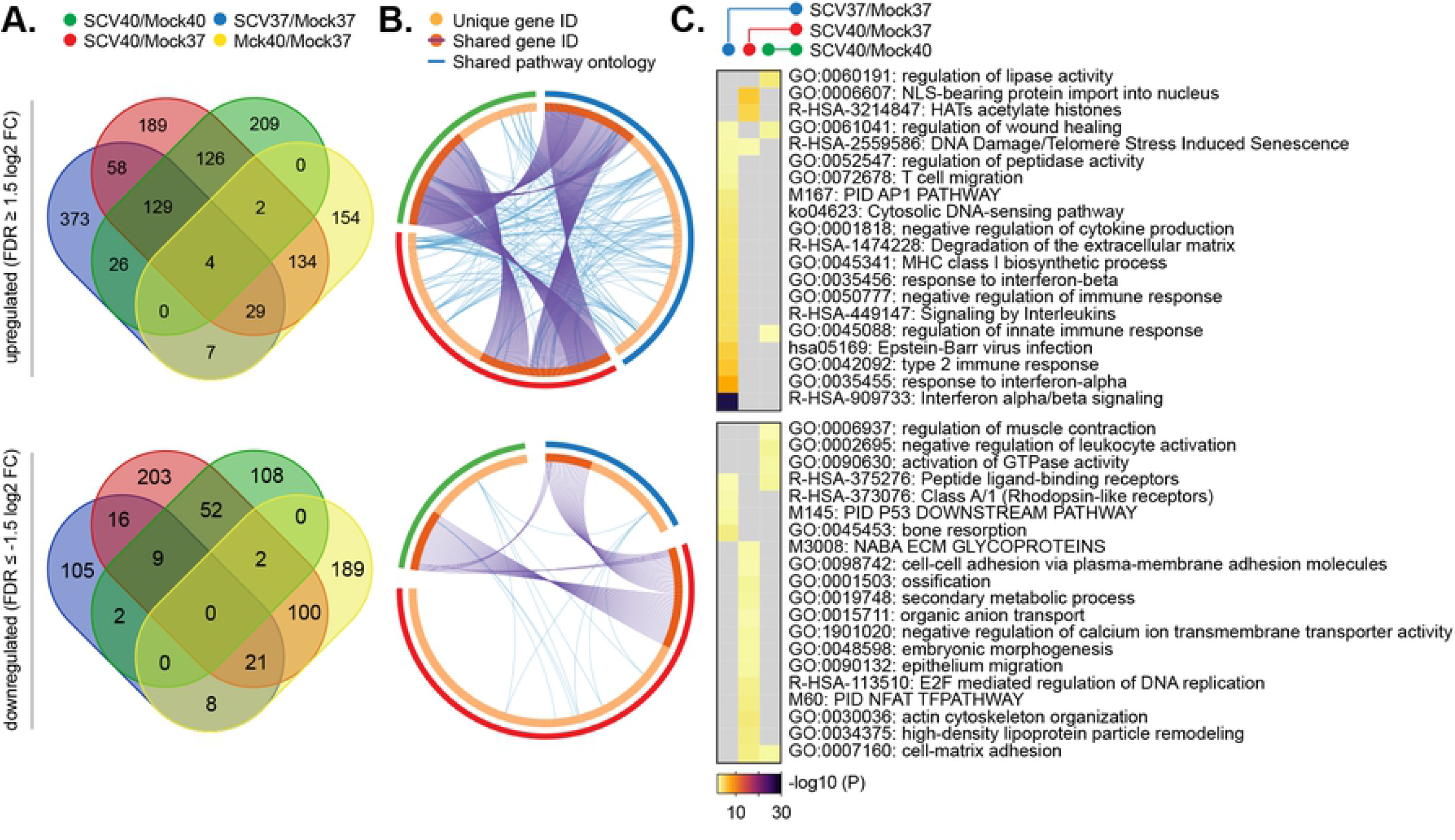
Respiratory airway cultures induce distinct transcriptional host responses to SARS-CoV-2 infection at elevated temperature. Ciliated respiratory cultures were incubated at 37 or 40°C for 24 h prior to mock (media only) or SARS-CoV-2 (SCV2; 10^4^ PFU/well) infection. Tissue were incubated at their respective temperatures for 72 h prior to RNA extraction and RNA-Seq. DEGs (p<0.05, ≥ 1.5 log2 FC [top panels] or ≤ -1.5 log2 FC [bottom panels]) were identified for each paired condition; blue ellipses, SCV2 37°C/Mock 37°C (SCV37/Mock37); green ellipses, SCV2 40°C/Mock 40°C (SCV40/Mock40); red ellipses, SCV2 40°C/Mock 37°C (SCV40/Mock37); yellow ellipses, Mock 40°C/Mock 37°C (Mock40/Mock37). (A) Venn diagram showing the number of unique or shared DEGs between each paired condition analyzed. (B) Circos plot showing the proportion of unique (light orange inner circle) or shared (dark orange inner circle + purple lines) DEGs between each paired condition analyzed. Blue lines, DEGs which share pathway gene ontology terms. (C) Metascape pathway analysis showing DEG enrichment p-value (-log10) for each paired condition analyzed. Grey boxes, p>0.05. (A to C) RNA-Seq data derived from RNA isolated from three technical replicates per sample condition. Data analysis is presented in Supplemental Files S1 to S5.

**Figure 6.**
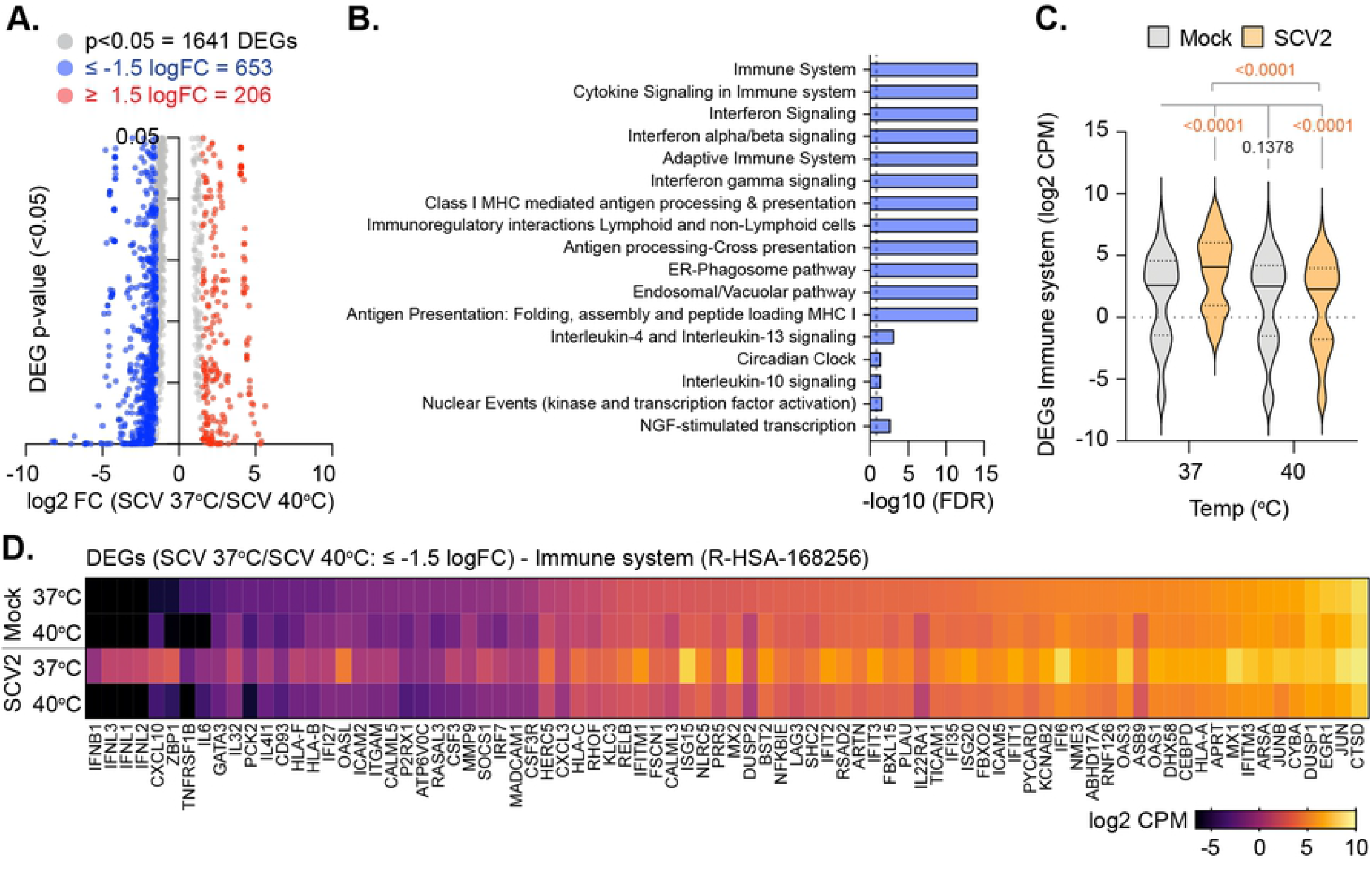
Elevated temperature restricts SARS-CoV-2 replication in respiratory airway cultures independently of the induction of IFN-mediated innate immune defences. Ciliated respiratory cultures were incubated at 37 or 40°C for 24h prior to mock (media) or SARS-CoV-2 (SCV2; 10^4^ PFU/well) infection. Tissue were incubated at their respective temperatures for 72 h prior to RNA extraction and RNA-Seq. (A) Scatter plots showing high confidence (p<0.05) differentially expressed gene (DEG) transcripts identified between 37 and 40°C SCV2 infected respiratory cultures (grey circles); DEGs ≥ 1.5 log2 FC, red circles; DEGs ≤ -1.5 log2 FC, blue circles. (B) Reactome pathway analysis of mapped DEGs (p<0.05, ≤ -1.5 log2 FC). Pathways enriched for DEGs with an FDR< 0.05 (plotted as -log10 FDR) shown (blue bars). Dotted line, threshold of significance (-log10 FDR of 0.05). (C) Expression profile (log2 CPM) of Immune system (R-HSA168256) DEGs (identified in B) relative to expression levels in Mock tissue incubated at 37 and 40^°^C. Black line, median; dotted lines, 5^th^ and 95^th^ percentile range; p-values shown, Wilcoxon matched-pairs signed rank test (top), one-way ANOVA Friedman test (bottom). (D) Expression levels (log2 CPM) for individual Immune system (R-HAS-168256) DEGs (identified in B) relative to mock at 37 and 40°C. (A to D) RNA-Seq data derived from RNA isolated from three technical replicates per sample condition. Data analysis is presented in Supplemental File S7.

In order to substantiate these findings, we examined the influence of temperature on SCV2 replication in VeroE6 cells, a cell line derived from *Chlorocebus Sabaeus* (African green monkey) known to be permissive to SCV2 infection but defective in type-I IFN-mediated immune defences (26, 36, 37). SCV2 replication was restricted at 40°C, but not 39°C, relative to infection at 37°C with or without elevated temperature pre-incubation (Figure 7A to C). Transcriptomic analysis revealed significant differences in the baseline expression of genes associated with the cellular response to heat pathway (GO:0034605) between respiratory tissue and undifferentiated HBE (p=0.0142) or VeroE6 (p<0.0001) cells (Figure 7D, E), but not a reference set of genes expressed across a wide range of cell types and tissues (Figure 7F, G, Supplemental File S8; (33-35)). We posit that such differences in the constitutive expression of heat stress response genes is likely to influence the relative restriction of SCV2 upon temperature elevation in a cell type, and potentially species-specific, dependent manner. These data support our tissue analysis (Figure 4 to 6), which demonstrates that the temperature-dependent restriction of SCV2 to occur independently of the induction of IFN-mediated antiviral immune defences.

**Figure 7.**
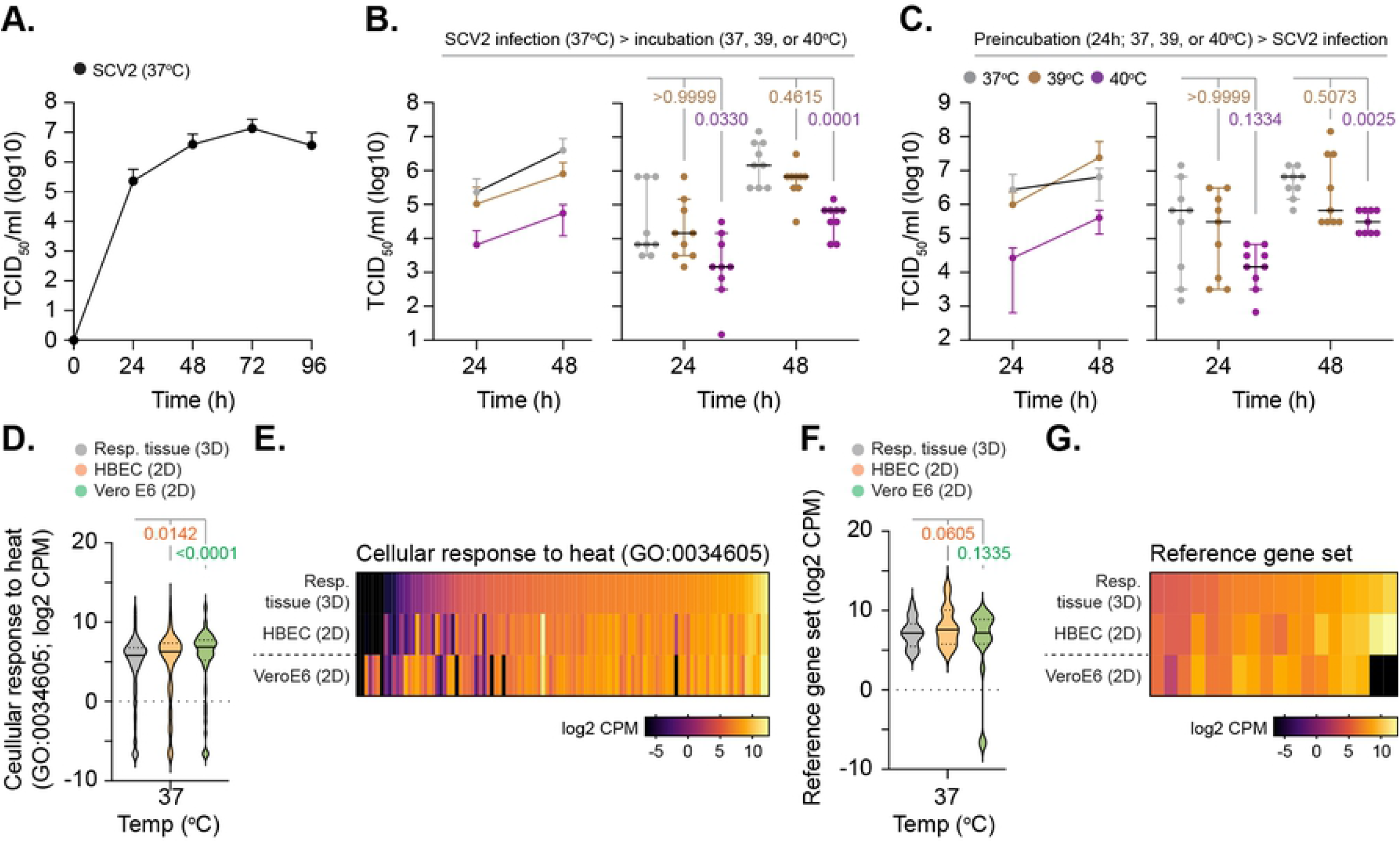
Elevated temperature restricts SARS-CoV-2 replication in VeroE6 cells. Permissive VeroE6 cells were infected with SARS-CoV-2 (SCV2; 10^4^ PFU/well) at 37°C prior to temperature elevation and incubation at 37, 39, or 40°C (A/B) or pre-incubated at 37, 39, or 40°C for 24 h prior to infection and continued incubation at their respective temperatures (C). (A) TCID_50_ growth curve of SCV2 infected VeroE6 cells incubated at 37°C over time (h). Means and SD shown. (B/C) TCID_50_ viral titres at 24 and 48 h post-infection. Left-hand panel; means and SD. Right-hand panel; black line, median; whisker, 95% confidence interval; all data points shown; p-values shown, one-way ANOVA Kruskal-Wallis test. (A-C) N=9 derived from a minimum of three independent biological experiments per sample condition. (D) Expression profile (log2 CPM) for genes associated with cellular response to heat (GO:0034605) in respiratory cultures (Resp. tissue), undifferentiated primary HBE, and VeroE6 cells incubated at 37°C; p-values shown, One-way ANOVA Friedman test. (E) Expression levels (log2 CPM) of individual genes associated with cellular response to heat (as in D). (F) Expression profile (log2 CPM) of a reference gene set (18 genes; (33-35)); p-values shown, One-way ANOVA Friedman test. (G) Expression values (log2 CPM) of a reference gene set (as in F). (D to G) RNA-Seq data derived from RNA isolated from three technical replicates per sample condition. Data analysis is presented in Supplemental File S8.

We next examined the stage at which SCV2 infection became restricted during infection of respiratory tissue at elevated temperature. RNA-Seq analysis of SCV2 infected tissues at 72 h post-infection demonstrated no significant difference in the total number of mapped reads (human + SCV2) between 37 and 40°C (Figure 8A, p=0.3429). However, analysis of viral reads identified a decrease in the relative abundance of viral transcripts mapping to multiple ORFs upon temperature elevation (Figure 8B to D, p<0.0010). Indirect immunofluorescence staining of tissue sections for SCV2 nucleocapsid (N) expression demonstrated significantly fewer clusters of SCV2 positive cells within infected tissues at 40°C relative to 37°C (Figure 8E, F, p=0.0022). We conclude that infection of respiratory tissue at elevated temperature restricts SCV2 replication through a mechanism that inhibits viral transcription independently of the induction of IFN-mediated antiviral immune defences that have been reported to restrict SCV2 propagation and spread (31, 38).

**Figure 8.**
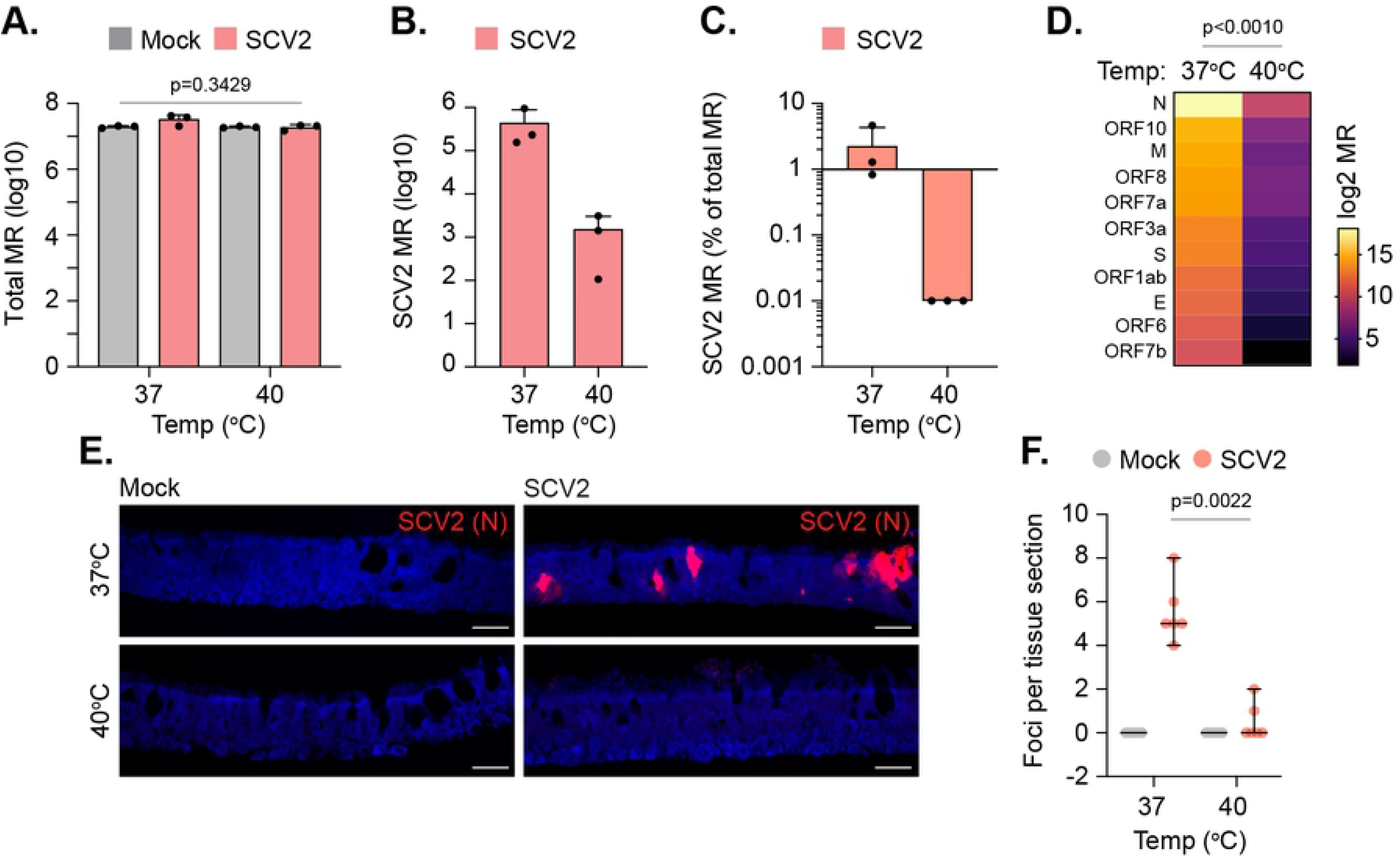
Elevated temperature restricts SARS-CoV-2 transcription in respiratory airway cultures. Ciliated respiratory cultures were incubated at 37 or 40°C for 24 h prior to mock (media only) or SARS-CoV-2 (SCV2; 10^4^ PFU/well) infection. Tissue were incubated at their respective temperatures for 72 h prior to RNA extraction and RNA-Seq. (A) Total mapped reads (MR; human + SCV2) from infected tissues; means and SD shown; p=0.3429, one-way ANOVA Kruskal-Wallis test. (B) SCV2 mapped reads from infected tissues; means and SD shown. (C) % SCV2 mapped reads of total mapped read count (human + SCV2); means and SD shown. (D) Expression values (log2 MR) of SCV2 gene transcripts; p<0.0010, Wilcoxon matched-pairs sign rank test. (A-D) RNA-Seq data derived from RNA isolated from three technical replicates per sample condition. (E) Indirect immunofluorescence staining of tissue sections showing SCV2 nucleocapsid (N, red) epithelial localization. Nuclei were stained with DAPI. Scale bars = 20 µm. (F) Quantitation of SCV2 N infectious foci in respiratory tissue sections. N=6 independently stained tissue sections per sample condition. Black line, median; whisker, 95% confidence interval; all data points shown; p=0.0022, Mann-Whitney *U*-test.

## Discussion

A defining symptom of COVID-19 is the onset of fever with a febrile temperature range of 38 to 41°C (4-8). However, the effect of elevated temperature on SCV2 tissue tropism and replication has remained to be determined. Here we identify a temperature-sensitive phenotype in SCV2 replication in respiratory epithelial tissue that occurs independently of the induction of IFN-mediated innate immune defences known to restrict SCV2 replication (31, 38). The differentiation of pseudostratified respiratory epithelium has proven to be a valuable research tool to investigate the cellular tropism, replication kinetics, and immune regulation of SCV2 infection, as these tissue models mimic many aspects of infection observed in animal models and COVID-19 patients (16-24, 29, 32, 39-42). While the use of such 3D models represents an important advancement over traditional 2D cell culture systems, the absence of circulating immune cells (e.g. macrophages, natural killer cells, dendritic cells, and neutrophils) which modulate fever and pro-inflammatory immune responses to microbial infection is an important limiting factor (9, 10, 43). Thus, we limit our conclusions to the effect of elevated temperature on SCV2 infection and replication within the respiratory epithelium. Additional animal studies are warranted to determine the overall net effect of fever and pro-inflammatory immune responses on body temperature thermoregulation and its corresponding influence on SCV2 tissue tropism and replication *in vivo*.

Consistent with previous reports (16, 18-21, 30-32), SCV2 infection of respiratory epithelium induced a pro-inflammatory immune response (Figure 2; IFNβ, IFNλ1-3, and IL-6). While SCV2 has been reported to induce a relatively weak IFN immune signature in comparison to other respiratory viruses (e.g. IAV) (20, 44), we regularly observed localized areas of epithelial infection (Figure 1A, 2E, 8E) that were coincident with elevated levels of ISG expression (Figure 2E, Mx1). Our findings are consistent with reports that have shown similar patterns of localized infection in the respiratory airway of SCV2 infected ferrets and surface epithelium of organoid cultures (29, 30, 32, 45). These data indicate that SCV2 induces a localized immune response during the opening phase of productive infection that is likely to restrict the progress of infection through the epithelium (Figure 1A) (30, 31, 38). Thus, differences in the proportion of epithelial infection and/or rate of intraepithelial spread may account for the observed differences in immune signatures between different respiratory pathogens (20, 46) and variance in apical yields of infectious virus at any given time point of analysis (Figure 1B and Figure 4B, D). As we observed localized areas of ISG induction (Figure 2E), our data support a model of epithelial infection where receptor-binding consumption kinetics of secreted IFN may limit cytokine diffusion and the intraepithelial induction of innate immune defences prior to the recruitment of immune cells and IFNγ secretion (47, 48).

We demonstrate that respiratory epithelial cultures induce a significant heat stress response upon temperature elevation without significant loss of tissue integrity, disruption to cellular transcription, diminished levels of ACE2 expression, or DAMP (damage associated molecular pattern) activation of immune defences at temperatures up to 40°C (Figure 3). Importantly, the elevation of temperature alone was not sufficient to inhibit SCV2 entry into respiratory tissue (Figure 4E), but restricted SCV2 replication leading to reduced levels of SCV2 apical shedding (Figure 4A to D). Thus, we identify tissue temperature to play an important role in the epithelial restriction of SCV2 replication. To our knowledge, these data represent the first account of a growth defect in SCV2 replication at febrile temperatures >37°C. Transcriptomic analysis identified substantial differences in the relative degree of immune activation in respiratory tissue infected at 40°C (Figure 5 and 6), despite these tissues having abundant levels of intracellular vRNA (Figure 4E). We posit that the lack of immune induction observed at 40°C is likely a consequence of diminished levels of SCV2 replication, as vRNA replication has previously been shown to play an important role in the production of PAMPs, including dsRNA intermediates, required for PRR detection and the activation of innate immune defences during respiratory virus infection (49-52). We also show the thermal restriction of SCV2 replication to occur in VeroE6 cells, a cell line permissive to SCV2 infection but defective in type-I IFN-mediated antiviral immune defences (26, 36, 37). Thus, we conclude that elevated temperature restricts SCV2 replication independently of the induction of ISG expression. Importantly, however, PRR detection of PAMPs which stimulate the production and secretion of pro-inflammatory cytokines (including IL-6) is a pre-requisite requirement for a fever response to microbial challenge *in vivo* (9, 10). Thus, additional animal studies are warranted to determine if the temperature restriction of SCV2 observed in respiratory cultures can occur independently of IFN-mediated immune defences *in vivo*. While speculative, we posit that low to moderate grade fever may confer protection to respiratory tissue within SCV2 infected individuals as a component of a homeostatically controlled non-hyperinflammatory immune response to infection.

While we identify temperature to play an important role in the regulation of SCV2 replication in respiratory epithelia, the precise mechanism(s) of restriction remains to be determined and warrants further investigation. The induction of a heat stress response alone within respiratory tissue may be sufficient to limit SCV2 replication (Figure 2, Figure S3), as the activation and/or induction of stress response proteins has been shown to play both a positive (proviral) and negative (antiviral) role in the replication of many viruses (53-55). Our transcriptomic analysis identified epithelial host responses to be differentially regulated in response to both temperature elevation and SCV2 infection (Figure 5 and 6), with unique profiles of induced gene expression between mock and SCV2 infected tissues at elevated temperature (Figure 5, Figure S2). Thus, SCV2 infection of respiratory epithelia at elevated temperature elicits a distinct host response that may directly contribute to the restriction of SCV2 propagation, for example the induction of mucins (Figure S2, *MUC4* and *MUC5AC*) which have been proposed to limit coronavirus disease progression and IAV replication (56, 57). We also identified a significant difference in the differential expression of lncRNAs and miRNAs upon temperature elevation and SCV2 infection that warrants additional investigation (Figure S4), as changes in the relative expression of non-coding RNAs are known to influence the outcome of virus infection independently of IFN-mediated immune defences (58). Thus, multiple gene products and/or pathways may contribute to the sequential or accumulative restriction of SCV2 replication within respiratory tissue at elevated temperature.

Importantly, we identify elevated temperature to lead to lower levels of SCV2 transcription and vRNA accumulation (Figure 4E, Figure 8D), suggesting that the temperature-dependent block in SCV2 propagation occurs prior to and/or during vRNA replication. We hypothesize that this block in transcription/replication may relate to the inhibition of SCV2 RNA-dependent RNA-polymerase (RdRp) activity or RdRp RNA binding affinity at elevated temperature, as the genomic replication activity of the IAV RdRp polymerase is known to be restricted at elevated temperature (41°C) (59). Notably, IAV cold-adaptation of RdRp activity has been shown to be a host determinant for avian IAV zoonosis, which naturally replicate at higher temperatures (41°C) within the intestinal tracts of birds (60). As circulating clinical strains of SCV2 are susceptible to non-synonymous mutations within RdRp coding sequences (http://cov-glue.cvr.gla.ac.uk/#/home), detailed molecular studies are warranted to determine if such amino acid substitutions influence the thermal restriction of SCV2 replication. To our knowledge, no temperature sensitive growth defect for coronaviruses has been reported to date, although a temperature-sensitive (ts) coronavirus mutant has been identified for murine hepatitis virus (MHV, tsNC11) (61). The replication defect of tsNC11 was attributed to coding substitutions within the macrodomain and papain-like protease 2 domain of the non-structural protein 3, which displayed a severe growth defect at 40°C in DBT (Delayed Brain Tumor) cells, whereas wild-type MHV replicated to equivalent titres at 40°C to those observed at 37°C (61). Thus, we present the first evidence demonstrating that a circulating clinical strain of coronavirus is sensitive to temperature thermoregulation. As such, heat-adaptation gain of function experiments through serial passage of SCV2 in respiratory cultures incubated at ≥39°C may shed light on whether the temperature restriction of SCV2 observed is related to viral or cellular host factors, although appropriate levels of biosafety (ethical and genetic modification) should be considered prior to such experimentation.

In summary, we identify an important role for tissue temperature in the restriction of SCV2 replication in respiratory epithelia that occurs independently of the induction of IFN-mediated antiviral immune defences. We demonstrate tissue temperature to significantly influence the differential regulation of epithelial host responses induced in response to SCV2 infection. Future investigation is warranted to determine the precise mechanism(s) of restriction, as this may uncover novel avenues for therapeutic intervention in the treatment of COVID-19.

## Materials and Methods

### Virus

Severe Acute Respiratory Syndrome Virus-2 (SARS-CoV-2, SCV2) BetaCoV/England/02/2020/EPI_ISL_407073 was isolated from a COVID-19 patient (a gift from Public Health England). The virus was passaged three times in VeroE6 cells and genotype sequence confirmed by Illumina sequencing. All experiments were performed in a Biosafety level 3-laboratory at the MRC-University of Glasgow Centre for Virus Research (SAPO/223/2017/1a).

### SCV2 replication kinetics in VeroE6 cells

VeroE6 cells (a gift from Michelle Bouloy, Institute Pasteur, France) were propagated in Dulbecco’s Modified Eagle Medium (DMEM GlutaMAX; ThermoFisher, 31966-021) supplemented with 10% fetal calf serum (FCS; ThermoFisher, 10499044 [Lot Number 08G8293K]) at 37°C with 5% CO_2_. Cells were seeded the day before infection at a density of 2×10^5^ cells per well in a 12-well plate and infected with 10^4^ PFU of SCV2 diluted in serum-free DMEM for 120 minutes (mins) with occasional rocking. The inoculum was removed and replaced by DMEM + 10% FCS. Supernatant was collected every 24 hours (h) and snap-frozen prior to analysis. For temperature inhibition experiments, cells were either pre-incubated at 37, 39, or 40°C for 24 h prior to infection with continued incubation at their respective temperatures or incubated at the indicated temperatures after the addition of the medium.

### Differentiation of respiratory epithelium

Primary human bronchiolar epithelial (HBE) cells from a healthy 63-year-old white Caucasian male (non-smoker) were sourced from Epithelix Sarl (Geneva, Switzerland). Cells were propagated at 37°C with 5% CO_2_ in hAEC culture medium (Epithelix, EP09AM). All differentiation experiments were performed on cells that had been passaged a total of four times. Cells were seeded onto transwells (Falcon, 734-0036) at a cell density of 3×10^4^ cells per transwell and grown to confluency. Cells were differentiated under air-liquid interface (ALI) in PneumaCult-ALI medium containing hydrocortisone and heparin supplements (STEMCELL Technologies, 05001) as specified by the manufacturer’s guidelines. Cells were differentiated for a minimum of 35 days, with media changed every 48 h and twice weekly apical washing in serum-free DMEM after day 20 at ALI. ALI cultures were apically washed twice before infection, 24 h prior to infection and immediately preceding infection. Cultures were inoculated apically with 100 µl of serum-free DMEM containing 10^4^ PFU of SCV2 (based on VeroE6 titres) for 120 mins. The inoculum was removed, and the apical surface washed once in serum-free DMEM (0 h time point). Unless stated otherwise, ALI cultures were incubated at 37°C with 5% CO_2_. For temperature inhibition experiments, ALI cultures were pre-incubated at 37, 39, or 40°C for 24 h prior to infection with continued incubation at their respective temperatures. Apical washing for 30 mins in 200 µl of serum-free DMEM was used to collect infectious virus. Supernatants were divided into two 100 µl aliquots; one for virus quantitation (TCID_50_) and the other for vRNA extraction in TRIzol (ThermoFisher, 15596026; 1 in 3 dilution). Samples were stored at -80^°^C until required. Tissues were fixed at the indicated time points in 8% formaldehyde (Fisher Scientific, F/1501/PB17) for 16 to 24 h prior to paraffin embedding and processing.

### TCID_50_

VeroE6 cells (subclone 6F5, MESO) were seeded the day before infection into 96-well-plates at a density of 1.1×10^5^ cells/well in DMEM + 10% FCS at 37°C with 5% CO_2_. Virus supernatants were serially diluted 1:10 in DMEM + 2% FCS in a total volume of 100 μl. Cells were infected in triplicate and incubated for three days at 37°C with 5% CO_2_. The inoculum was removed and fresh media overlayed. Plates were incubated for three days at 37°C in 5% CO_2_. Cells were fixed in 8% formaldehyde (Fisher Scientific, F/1501/PB17) for ≥ 2.5 h. Cytopathic effect was scored by staining with 0.1% Coomassie Brilliant Blue (BioRad, 1610406) in 45% methanol and 10% glacial acetic acid. TCID_50_ was calculated according to Reed–Muench and Spearman–Kärber as described (62).

### Plaque assay

VeroE6 cells were seeded at a density of 2.5×10^5^ cells/well in 12-well plates the day before infection. Ten-fold serial dilutions of virus were prepared in DMEM + 2% FCS in a total volume of 500 μl. Cells were infected for 60 mins with occasional rocking at 37°C in 5% CO_2_. The inoculum was removed and 1 ml of overlay containing 0.6% Avicel (Avicel microcrystalline cellulose, RC-591) made up in DMEM containing 2% FCS was added to each well. Cells were incubated for three days at 37°C in 5% CO_2_ prior to fixation in 8% formaldehyde and Coomassie brilliant blue staining (as described above). Plaques were counted manually and the plaque forming unit (PFU) per ml calculated.

### Reverse transcription quantitative PCR (RTq-PCR)

RNA was extracted from TRIzol treated apical wash samples using a QIAGEN RNeasy Mini kit (Qiagen, 74106) following the manufacture’s protocol. SCV2 RNA was quantified using a NEB Luna Universal Probe One-Step RT-qPCR Kit (New England Biolabs, E3006) and 2019-nCoV CDC N1 primers and probes (IDT, 10006713). Genome copy numbers were quantified using a standard curve generated from serial dilutions of an RNA standard used throughout the study. The RNA standard was calibrated using a plasmid (2019-nCoV_N; IDT, 10006625) that was quantified using droplet digital PCR. Values were normalized to copies/ml for apical washes or copies/tissue for cell-associated RNA.

### *In situ-*hybridization and immunostaining of respiratory tissue sections

Paraffin-embedded tissues were sectioned using a microtome (∼2-3 µm thick) and mounted on glass slides. Tissue sections were stained with Haematoxylin and Eosin (H&E). For immunofluorescence staining, tissue sections underwent antigen retrieval by citrate (pH 6) or EDTA (pH 8) pressure cooking (as indicated). RNAscope was used for the detection of SCV2 RNA using a Spike-specific probe set (Advanced Cell Diagnostics, 848561 and 322372) following the manufacturer’s protocol, which included a pre-treatment with boiling in target solution and proteinase K treatment. ACE2 was detected using a rabbit polyclonal anti-hACE2 antibody (Cell Signalling, 4355S; citrate antigen-retrieval) and EnVision+ anti-rabbit HRP (Agilent, K4003). Slides were scanned with a bright field slide scanner (Leica, Aperio Versa 8). Mx1 was detected using a mouse monoclonal anti-Mx1 antibody ((63); EDTA antigen-retrieval). SCV2 nucleocapsid protein was detected using a sheep polyclonal anti-SCV2 N protein antibody (University of Dundee, DA114; EDTA antigen-retrieval). Secondary antibodies for detection were rabbit anti-sheep AlexaFluor 555 (abcam, ab150182) and rabbit anti-mouse AlexaFluor 488 (Sigma-Aldrich, SAB4600056). Nuclei were stained with ProLong Gold (Life Technologies, P36941). Images were collected using a Zeiss LSM 710 confocal microscope using 40x Plan-Apochromat oil objective lens (numerical aperture 1.4) using 405, 488, and 543 nm laser lines. Zen black software (Zeiss) was used for image capture and exporting images, with minimal adjustment (image rotation) in Adobe Photoshop for presentation.

### Tissue section image analysis

Scanned Haematoxylin and Eosin (H&E) stained sections were analyzed using Aperio ImageScope analysis software (Leica). Epithelial thickness was measured by manually outlining the bottom and top surfaces of each epithelium (excluding cilia) and measuring the vertical point-to-point distance between each surface across the epithelium. One hundred evenly distributed distance measurements were acquired per epithelium per experimental condition in triplicate.

### RNA sequencing (RNA-Seq)

Transwell tissue or undifferentiated cells seeded into 12-well dishes at a cell density of 2×10^5^ cells per well were harvested by scraping into TRIzol Reagent and transferred into tubes containing 2.8 mm ceramic beads (Stretton Scientific, P000916-LYSK0-A.0). Samples were homogenized (two 20 sec pulses with a 30 sec interval at room temperature, RT) using a Percellys Cryolys Evolution Super Homogeniser (Bertin Instruments, P000671-CLYS2-A) at 5500 rounds per minute. The homogenised suspension was loaded into QIAshredder tubes (Qiagen, 79654) and centrifuged (12,000 x g for 2 mins at RT). 0.25 ml of chloroform (VWR Life Sciences, 0757) was added to the eluate and incubated at RT for 7 mins prior to centrifugation (12’000 x g, 15 minutes, 4°C). The aqueous phase was transferred into a fresh tube, mixed with 250 μl of 100% ethanol, and RNA isolated using RNAeasy columns (Qiagen, 74104) following the manufacturer’s protocol, that included a 15 min DNAse treatment (Qiagen, 79254) treatment at RT. Eluted RNA was quantified using a NanoDrop 2000 Spectrophotometer (ThermoFisher Scientific, ND-2000) and quality controlled on a TapeStation (Agilent Technologies, G2991AA). All samples had a RIN score of ≥ 9. One microgram of total RNA was used to prepare libraries for sequencing using an Illumina TruSeq Stranded mRNA HT kit (Illumina, 20020594) and SuperScript2 Reverse Transcriptase (Invitrogen, 18064014) according to the manufacturer’s instructions. Libraries were pooled in equimolar concentrations and sequenced using an Illumina NextSeq 500 sequencer (Illumina, FC-404-2005). At least 95% of the reads generated presented a Q score of ≥ 30. RNA-Seq reads were quality assessed (FastQC; http://www.bioinformatics.babraham.ac.uk/projects/fastqc) and sequence adaptors removed (TrimGalore; https://www.bioinformatics.babraham.ac.uk/projects/trim_galore/). RNA-Seq reads were aligned to the *Homo sapiens* genome (GRCh38) or *Chlorocebus Sabaeus* genome (ChlSab1.1), downloaded via Ensembl using HISAT2. HISAT2 is a fast and sensitive splice aware mapper, which aligns RNA sequencing reads to mammalian-sized genomes using FM index strategy (64). RNA-Seq reads were also mapped to SARS-CoV-2 (GISAID accession ID:EPI_ISL_407073) using Bowtie2 (65). FeatureCount (66) was used to count reads mapping to gene annotation files. Reads counts were normalized to counts per million (CPM) unless otherwise stated. The edgeR package was used to calculate the gene expression level and to analyze differentially expressed genes between sample groups (67). Sequences have been deposited in the European Nucleotide Archive (https://www.ebi.ac.uk/ena/browser/home), accession number PRJEB41332. Only high confidence (p<0.05) differentially expressed genes (DEGs; ≥ 1.5 or ≤ -1.5 log2 fold change, log2 FC) were used for pathway analysis in Reactome (https://reactome.org) (68, 69) or differential pathway enrichment in Metascape (https://metascape.org/gp/index.html#/main/step1) (70). In Reactome, the gene mapping tool was used as a filter to identify pathways enriched (over-represented) for mapped DEG entities. FDR (False Discovery Rate) values <0.05 were considered significant for pathway enrichment. In Metascape, all DEGs were used for differential pathway analysis. Pathway p-values <0.05 were considered significant. Differential expressed (p<0.05, ≥ 1.5 or ≤ -1.5 log2 FC) lncRNAs and miRNAs were identified using the Ensembl BioMart tool (http://www.ensembl.org/biomart/martview/05285d5f063a05a82b8ba71fe18a0f18). Heat maps were plotted in GraphPad Prism (version 9). Mean counts per million (CPM) values of zero were normalized to 0.01 for log2 presentation. Venn diagrams were plotted using http://bioinformatics.psb.ugent.be/webtools/Venn/.

### Statistical analysis

GraphPad Prism (version 9) was used for statistical analysis. For unpaired non-parametric data, a Kruskal-Wallis one-way ANOVA or Mann-Whitney *U*-test was applied. For paired non-parametric data, a one-way ANOVA Friedman test or Wilcoxon matched-pairs sign rank test was applied. Statistical p-values are shown throughout. Statistically significant differences were accepted at p≤0.05.

## Acknowledgements

The authors thank Lynn Marion Stevenson, Frazer Bell, and Lynn Oxford (College of Medical, Veterinary and Life Sciences, University of Glasgow) for their exceptional efforts during the UK lockdown. VH was funded by the German Research Foundation (Deutsche Forschungsgemeinschaft; project number 406109949) and the Federal Ministry of Food and Agriculture (BMEL; Förderkennzeichen: 01KI1723G). KD and PRM were funded by the Medical Research Council (MRC; MC_UU_12014/9 to PRM). JW was funded by an MRC CVR DTA award (MC_ST_U18018). DG was funded by an MRC-DTP award (MR/R502327/1). CR was funded by a BBSRC-CTP award (BB/R505341/1). QG was funded by the MRC (MC_UU_12014/12). IE was funded by a CSO project grant (TCS/19/11). MES was funded by the MRC (MC PC 19026). AMS was funded by a UKRI/DHSC grant (BB/R019843/1 to Brian Willett, MRC-UoG CVR) and MRC CoV supplement grant (MC_PC_19026). RMP was funded by the MRC (MC_UU_12014/10). AMG was funded by studentship awards from the University of Glasgow School of Veterinary Medicine (Georgina D. Gardner, 145813; John Crawford, 123939). CB and SMF were funded by the MRC (MC_UU_12014/5 to CB).

## Availability of data

The datasets generated and analyzed during the current study are available in the University of Glasgow data repository (a doi will be generated upon manuscript acceptance). RNA-Seq data sets are available from the European Nucleotide Archive, accession number PRJEB41332 (available upon manuscript acceptance).

## Author contributions

VH, KD (joint first authors): Conceptualization, Data curation, Formal Analysis, Investigation, Methodology, Project administration, Validation, Visualization, Writing – original draft, Writing – review & editing.

JKW, RJ, CR, DG, IE, AS, SMF, QG: Data curation, Formal Analysis, Investigation, Methodology, Validation

MS, AMS, RMP.: Resources, Methodology

AMG: Data curation, Formal Analysis

SG, PRM: Conceptualization, Funding acquisition, Methodology, Supervision, Writing – review & editing.

CB: Conceptualization, Methodology, Data curation, Formal Analysis, Investigation, Funding acquisition, Project administration, Supervision, Validation, Writing – Original draft, Writing – review & editing.

## Figure legends

**Figure S1. Respiratory airway cultures induce a heat stress response**. Ciliated respiratory cultures were incubated at 37 or 40°C for 72 h prior to RNA extraction and RNA-Seq. Heat map showing expression values (log2 CPM) of genes associated with the cellular response to heat pathway (GO: 0034605); p=0.0018, Wilcoxon matched-pairs sign rank test; every second row labelled. RNA-Seq data derived from RNA isolated from three technical replicates per sample condition. Data analysis is presented in Supplemental File S2.

**Figure S2. Identification of DEGs in mock and SARS-CoV-2 infected respiratory cultures incubated at 40°C**. Ciliated respiratory cultures were incubated at 40°C for 24 h prior to mock (media only) or SARS-CoV-2 (SCV2; 10^4^ PFU/well) infection and continued incubation at 40°C. Tissues were harvested at 72 h for RNA extraction and RNA-Seq. (A) Scatter plots showing high confidence (p<0.05) differentially expressed gene (DEG) transcripts identified between mock and SCV2 infected respiratory cultures at 40°C (grey circles); DEGs ≥ 1.5 log2 FC, red circles; DEGs ≤ -1.5 log2 FC, blue circles. (B) Reactome pathway analysis of mapped DEGs (p<0.05, ≥ 1.5 log2 FC [red bars] or ≤ -1.5 log2 FC [blue bars]). Top 10 enriched pathways shown. Dotted line, threshold of significance (-log10 FDR of 0.05). (C) Expression values (log2 CPM) of Reactome mapped DEGs; p-values shown, Wilcoxon matched-pairs sign rank test. (A to C) RNA-Seq data derived from RNA isolated from three technical replicates per sample condition. Data analysis is presented in Supplemental File S3.

**Figure S3. Identification of DEGs in mock and SARS-CoV-2 infected respiratory cultures incubated at 37 and 40°C, respectively**. Ciliated respiratory cultures were incubated at 37 or 40°C for 24 h prior to mock (media only) or SARS-CoV-2 (SCV2; 10^4^ PFU/well) infection and continued incubation at their respective temperatures. Tissues were harvested at 72 h for RNA extraction and RNA-Seq. (A) Scatter plots showing high confidence (p<0.05) differentially expressed gene (DEG) transcripts identified between mock (37°C) and SCV2 infected (40°C) respiratory cultures (grey circles); DEGs ≥ 1.5 log2 FC, red circles; DEGs ≤ -1.5 log2 FC, blue circles. (B) Reactome pathway analysis of mapped DEGs (p<0.05, ≥ 1.5 log2 FC [red bars, every sixth row labelled] or ≤ -1.5 log2 FC [blue bars]). Dotted line, threshold of significance (-log10 FDR of 0.05). Black arrow, identification of cellular response to heat stress pathway. (C) Expression values (log2 CPM) of Reactome mapped DEGs; p-values shown, Wilcoxon matched-pairs sign rank test. Every third column labelled. (A to C) RNA-Seq data derived from RNA isolated from three technical replicates per sample condition. Data analysis is presented in Supplemental File S4.

**Figure S4. Respiratory airway cultures induce distinct lncRNA and miRNA transcriptional host responses to SARS-CoV-2 infection at elevated temperature**. Ciliated respiratory cultures were incubated at 37 or 40°C for 24 h prior to mock (media only) or SARS-CoV-2 (SCV2; 10^4^ PFU/well) infection. Tissue were incubated at their respective temperatures for 72 h prior to RNA extraction and RNA-Seq. DEGs (p<0.05, ≥ 1.5 log2 FC [top panels] or ≤ -1.5 log2 FC [bottom panels]) were identified for each paired condition; blue circles/ellipses, SCV2 37°C/Mock 37°C (SCV37/Mock37); green circles/ellipses, SCV2 40°C/Mock 40°C (SCV40/Mock40); red circles/ellipses, SCV2 40°C/Mock 37°C (SCV40/Mock37); yellow circles/ellipses, Mock 40°C/Mock 37°C (Mock40/Mock37). (A) Proportion of lncRNA + miRNA (grey numbers and lines [relative %]) identified per DEG population (coloured numbers and lines [relative %]). (B) Venn diagram showing the number of unique or shared lncRNA + miRNAs identified between each paired condition analyzed. (C) Circos plot showing the proportion of unique (light orange inner circle) or shared (dark orange inner circle + purple lines) lncRNA + miRNAs between each paired condition analyzed. (D) Expression values (log2 CPM) of lncRNA + miRNA identified per sample condition analyzed; p-values shown, one-way ANOVA Friedman test (top), Wilcoxon matched-pairs sign rank test (bottom). (A to D) RNA-Seq data derived from RNA isolated from three technical replicates per sample condition. Data analysis is presented in Supplemental File S6.

## Supplementary Files

**Supplemental File S1**. RNA-Seq data analysis for mock (Mck)-treated or SARS-CoV-2 (SCV2) infected respiratory tissues at 37°C.

**Supplemental File S2**. RNA-Seq data analysis for mock (Mck)-treated respiratory tissues incubated at 37 and 40°C.

**Supplemental File S3**. RNA-Seq data analysis for mock (Mck)-treated or SARS-CoV-2 (SCV2) infected respiratory tissues at 40°C.

**Supplemental File S4**. RNA-Seq data analysis for mock (Mck)-treated or SARS-CoV-2 (SCV2) infected respiratory tissues at 37°C and 40°C, respectively.

**Supplemental File S5**. Comparative DEG analysis of RNA-Seq data derived from mock (Mck)-treated or SARS-CoV-2 (SCV2) infected respiratory tissues at 37 and 40°C. Supplemental Files S1-S4.

**Supplemental File S6**. Comparative lncRNA and miRNA DEG analysis of RNA-Seq data derived from mock (Mck)-treated or SARS-CoV-2 (SCV2) infected respiratory tissues at 37 and 40°C. Supplemental Files S1-S4.

**Supplemental File S7**. RNA-Seq data analysis for SARS-CoV-2 (SCV2) infected respiratory tissues at 37 and 40°C.

**Supplemental File S8**. DEG analysis of the Cellular response to heat (GO:0034605) from RNA-Seq data derived from mock (Mck)-treated respiratory tissues (Resp. tissue) and undifferentiated HBEC and VeroE6 cells incubated at 37°C.

